# Oestrogenic vascular effects are diminished by ageing

**DOI:** 10.1101/108274

**Authors:** Christopher J. Nicholson, Michèle Sweeney, Stephen C. Robson, Michael J. Taggart

## Abstract

The beneficial role of oestrogen in the vascular system may be due, in part, through reduction of peripheral vascular resistance. The use of oestrogen therapy to prevent cardiovascular disease in post-menopausal women remains contentious. This study investigated the influence of the menopause and ageing on the acute vasodilatory effects of oestrogen in *ex vivo* uterine resistance arteries.

Vessels were obtained from young (2.9 ± 0.1 months) and aged (24.2 ± 0.1 and 28.9 ± 0.3 months) female mice and pre-(42.3 ± 0.5 years) and post-menopausal (61.9 ± 0.9 years) women. Ageing was associated with profound structural alterations of murine uterine arteries, including the occurrence of outward hypertrophic remodelling and increased stiffness. Endothelial and smooth muscle function were diminished in uterine (and tail) arteries from aged mice and post-menopausal women. The acute vasodilatory effects of 17β-oestradiol (non-specific oestrogen receptor (ER) agonist), PPT (ERα-specific agonist) and DPN (ERβ-specific agonist) on resistance arteries were attenuated by ageing and the menopause. However, the impairment of oestrogenic relaxation was evident after the occurrence of age-related endothelial dysfunction and diminished distensibility. The data indicate, therefore, that adverse resistance arterial ageing is a dominant factor leading to weakened vasodilatory action of oestrogenic compounds.

## Introduction

Cardiovascular diseases (CVDs) are the leading cause of death worldwide (Mozaffarian et al., 2015). Age is the biggest determinant of an individual’s cardiovascular health and since the ageing population is growing substantially, there is a need to develop a greater understanding of the influence of age on the cardiovascular system (North and Sinclair, 2012). There are also marked differences in the prevalence of CVD in men and women; prior to 50 years of age, the average age of menopause onset, the risk is considerably lower in women but thereafter the incidence of CVDs is similar (Leening et al., 2014; Lerner and Kannel, 1986). Post-menopausal deprivation of female sex hormones, primarily oestrogen, may be related to the increased CVD risk in ageing women (Kuhl, 2005).

Oestrogen binds to two receptors, ERα and ERβ, which can then initiate ligand-activated transcription factor signalling (Miller and Duckles, 2008). However, oestrogen can also induce acute arterial vasodilation separate from genomic-directed actions (Al Zubair et al., 2005; Bolego et al., 2005; Corcoran et al., 2014; Montgomery et al., 2003; Patkar et al., 2011; Scott et al., 2007; Zhou et al., 2013). It has been proposed that oestrogen activates endothelial nitric oxide synthase (and thus NO production), endothelial prostacyclin and endothelium-derived hyperpolarizing (EDH) factor, which act on adjacent smooth muscle cells to induce relaxation (Al Zubair et al., 2005; Corcoran et al., 2014; Zhou et al., 2013). Indeed, it has been suggested that the contribution of EDH is greater in females due to the presence of oestrogen (Kong et al., 2015; Leung and Vanhoutte, 2017; Wong et al., 2014). Further to acting through the classical ERs, oestrogen has recently been demonstrated to act through the unrelated G-protein-coupled oestrogen receptor (GPER1) to induce vasodilation (Arefin et al., 2014; Broughton et al., 2010; Haas et al., 2009; Lindsey et al., 2009; Lindsey et al., 2013). In large arteries, the effects of oestrogen are considered to protect against inflammatory-related disorders such as atherosclerosis, whereas in the resistance vasculature, the acute vasodilatory actions may lower peripheral resistance and protect against hypertension (Kublickiene et al., 2005).

Consistent with the proposed cardiovascular protective role of oestrogen, early observational studies, such as the Nurses’ Health Study, suggested oestrogen therapy might reduce the risk of CVD in post-menopausal women (Grodstein et al., 2000). However, randomised clinical trials largely failed to show any cardiovascular benefits of menopausal hormone therapy (MHT) in post-menopausal women, with guidelines concluding that MHT should not be used for CVD prevention (Duvernoy and Mosca, 2002; Grady et al., 2002; Rossouw et al., 2002). However, subsequent re-analysis of the Women’s Health Initiative data revealed a trend for a lower risk of coronary heart disease in hormone-treated women from the youngest age group. This suggested the possibility of a ‘timing’ hypothesis wherein MHT initiated within 10 years of the menopause could be beneficial to CVD health (Grodstein et al., 2006; Rossouw et al., 2007; Valdiviezo et al., 2013). Indeed, a recent study suggested there to be some beneficial cardiovascular effects of oestradiol treatment in early (< 6 years) menopausal compared to late (≥ 10 years) menopausal women (Hodis et al., 2016). Recent evidence has also highlighted the role for ageing per se, independent of menopausal status, to cardiovascular risk in women (de Kat et al., 2017; Vaidya et al., 2011). Thus, it remains an important objective to elucidate the relative contributions of ageing and the menopause, and the influence of oestrogenic compounds, on vascular function.

Therefore, in the present study we have sought to determine the contributions of age or the menopause on arterial function and oestrogenic responsiveness *in vitro*. To do so we employed two experimental models of arterial vascular function; uterine and tail arteries from young (3 months) and old (24 and 29 months) mice, to determine the influence of age, and uterine arteries from non-pregnant women ranging in age from 32-75 years, to assess the influence of menopause and/or age.

## Results

### Ageing negatively regulates murine uterine arterial function and passive structural characteristics

Prior to testing the influence of age on oestrogenic responsiveness of murine resistance arteries, we sought to determine the role of ageing on vasoactive and structural properties of these vessels. We utilised mice from the C57BL background (C57BL/Ircfa) from 3 separate age groups (Table 1). These mice remain generally healthy into old age with no strain-specific pathological conditions (Greaves et al., 2011; Lister et al., 2010; Martin et al., 1998; Rowlatt et al., 1976). Vasoactive responses to the thromboxane agonist U46619 (U4), the endothelium-dependent vasodilator acetylcholine (ACh) and the passive structural responses to increasing intravascular pressure (illustrated in Figure 1) were assessed. Both U4-induced constriction (maximum contractions for 3-month-old: 20.7 ± 0.8 kPa, 24-month-old: 16.7 ± 0.9 kPa and 29-month-old: 15.4 ± 1.2 kPa, Figure 1A) and endothelium-dependent vasodilation (maximum relaxations for 3-month-old: 55.1 ± 1.8%, 24-month-old: 42.1 ± 2.4% and 29-month-old: 42.9 ± 3.6%, Figure 1B) were attenuated in aged mice.

**Table 1:** Mouse details. All values, apart from oestrous stage, are presented as mean ± SEM. * P < 0.05 from young mice (ordinary one-way ANOVA).

| Characteristics | 3 months (n=26) | 24 months (n=10) | 29 months (n=5) |
| --- | --- | --- | --- |
| **Age (months)** | 2.91 ± 0.09 | 24.18 ± 0.14* | 28.86 ± 0.28* |
| **Weight (g)** | 22.06 ± 0.08 | 26.83 ± 0.44* | 30.48 ± 1.48* |
| **Uterine weight (g)** | 0.46 ± 0.02 | 0.67 ± 0.05 | 0.34 ± 0.06 |
| **Heart weight (g)** | 0.16 ± 0.01 | 0.16 ± 0.00 | 0.16 ± 0.01 |
| **Oestrous stage:** |  |  |  |
| Pro-oestrus | 9 | - | - |
| Oestrus | 6 | - | - |
| Metoestrus | 4 | - | - |
| Dioestrus | 7 | 10 | 5 |
| **Resting diameter (µm):** |  |  |  |
| Uterine arteries | 164 ± 4.5 | 188.5 ± 4.0* | 211.0 ± 5.0* |
| Tail arteries | 251.7 ± 6.3 | 227.0 ± 5.5 | 265.9 ± 9.7 |

**Figure 1:**
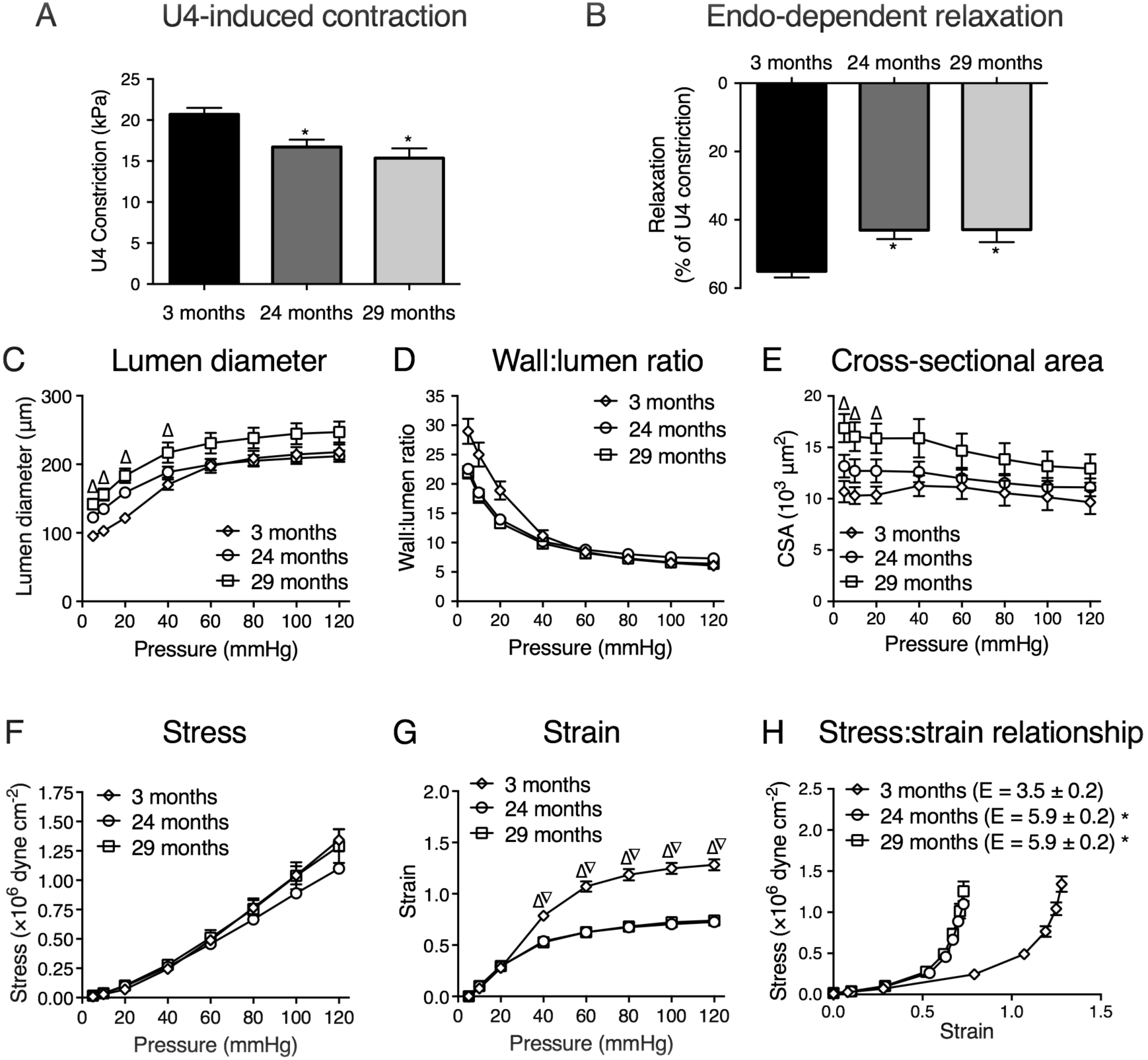
Ageing alters the vasoactive and passive structural characteristics of murine uterine arteries. For the vasoactive experiments, uterine arteries from 3- (n=26), 24- (n=10), and 29-month-old (n=5) mice were mounted on a wire myograph and exposed to the thromboxane agonist U4 (10^−6^ M, A). Following steady-state constriction, endothelium-dependent relaxation was assessed by the addition of acetylcholine (10^−5^M, B). * P < 0.05 from young mice (ordinary one-way ANOVA). For the assessment of passive structural characteristics, separate uterine arteries from 3- (n=5), 24- (n=7), and 29-month-old (n=4) mice were mounted on a pressure myograph and measurements of (C) vessel diameter and wall thickness were used to calculate (D) wall: lumen ratio, (E) CSA, (F) vessel stress and (G) strain at each pressure step (0 - 120 mmHg) in calcium-free conditions. P < 0.05 from 24-month-old mice (∇) or 30-month-old mice (∆) (repeated measures two-way ANOVA). Vessel stress was plotted against vessel strain to assess the stress: strain relationship, which is used to ascertain arterial distensibility (H). E = Elastic modulus. * P < 0.05 from young mice (unpaired *t-test*). Data are presented as mean ± SEM.

The effect of ageing on passive structural characteristics was more complex. There was no difference in the passive structural characteristics of uterine arteries from 3-and 24-month-old mice. However, further ageing to 29 months increased the passive diameter of uterine arteries (Figure 1C), which was associated with an increase in cross-sectional area (CSA) (Figure 1E). There was also a slight, but non-statistically significant, decrease in the wall-to-lumen ratio (Figure 1D). Increases in passive diameter and CSA are consistent with outward hypertrophic remodelling. Ageing was also associated with a decrease in the strain: pressure relationship (Figure 1G) with no change in the stress: pressure relationship (Figure 1F). Consequently, there was an increased elastic modulus in arteries from aged mice (Figure 1H), indicating a reduced arterial distensibility (i.e. increased stiffness). This is supported by a leftward shift in the stress: strain curve in uterine arteries from aged mice (Figure 1H).

### Oestrogenic responsiveness of murine uterine arteries is impaired in the oldest mice, after the occurrence of endothelial dysfunction and increased stiffness

We examined the effect of ageing on the vasodilatory responses of resistance arteries to oestrogenic compounds. Figure 2 demonstrates the vasorelaxant effects of 17β-oestradiol (17β), PPT (ERα) and DPN (ERβ) on pre-constricted uterine arteries from 3-, 24- and 29-month-old mice. Ageing was associated with attenuation of the acute vasodilatory properties of all three oestrogenic compounds but the effects were agonist-specific. The acute vasodilatory effects of 17β and PPT, which induced the strongest responses in these vessels, were not different between 3 and 24 months but were reduced when comparing 3-and 30-month-old mice. Therefore, impaired relaxations to oestrogenic compounds occurred after altered functional and elastic arterial properties of these vessels were demonstrable (see Figure 1).

**Figure 2:**
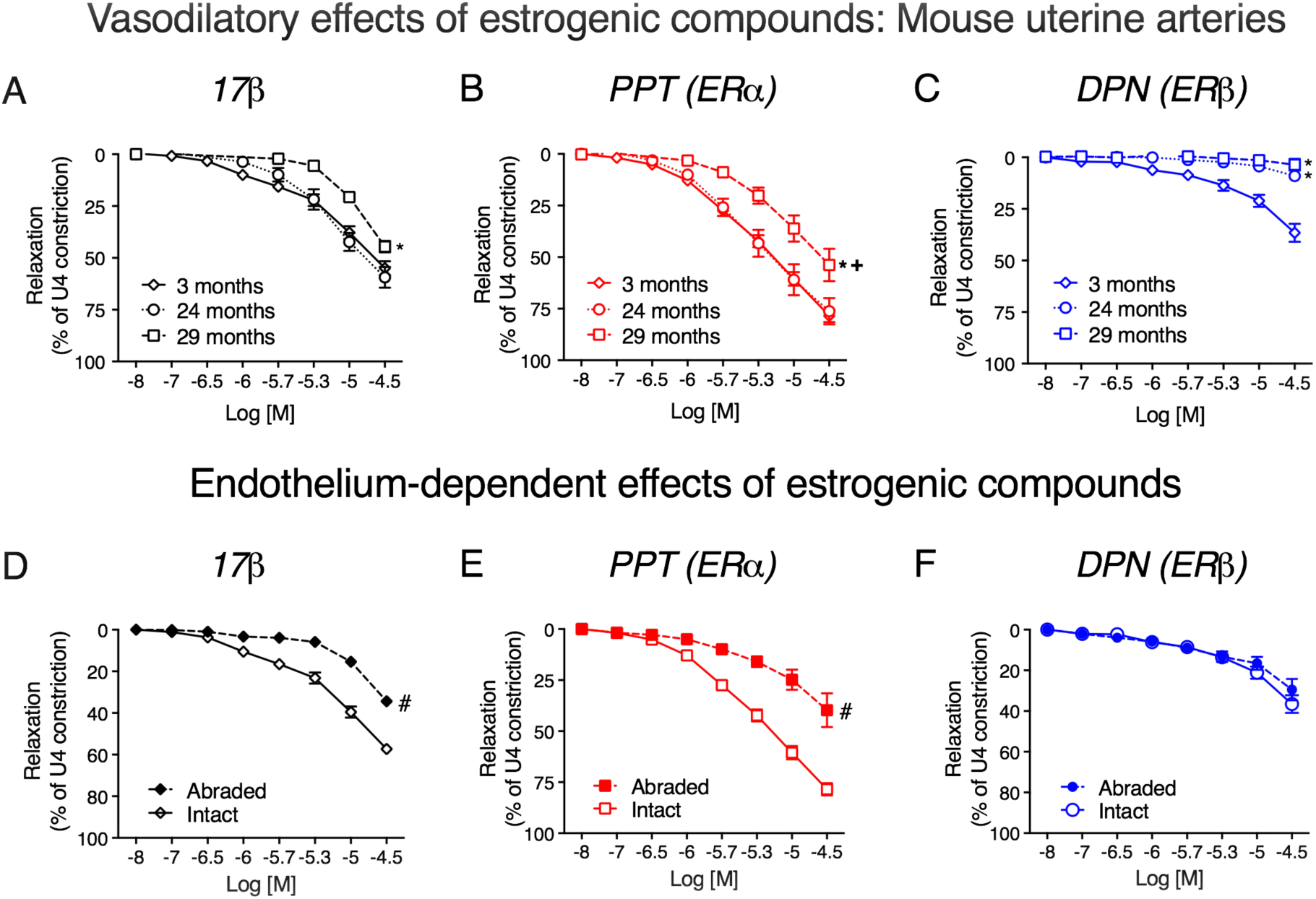
The acute responses of murine uterine arteries to oestrogenic compounds are attenuated by ageing. Acute relaxations to 17β (A), PPT (B) and DPN (C) were compared in uterine arteries from 3- (n=26), 24- (n=9) and 29-month-old (n=5) female mice. P < 0.05 from young mice (*) or 24-month-old mice (+) (repeated measures two-way ANOVA). In a separate set of experiments, pre-constricted (U4, 10^−6^M) endothelium-intact (solid lines and open symbols, n=26) and endothelium-abraded (dashed lines and closed symbols, n=7) uterine arteries from young mice were evaluated for their responses to incremental doses (10^−8^ M – 10^−4.5^ M) of 17β-oestradiol (D), PPT (E) or DPN (F). # P < 0.05 vs. intact arteries (repeated measures two-way ANOVA). Data are presented as mean ± SEM.

Having determined that endothelial dysfunction was associated with ageing in murine resistance arteries, we hypothesised that the reduced vasorelaxant effects of oestrogenic compounds may be attributable, in part, to altered endothelial-mediated relaxation. We therefore examined the relaxatory effects of each oestrogenic compound in endothelium-intact and-abraded uterine vessels (demonstrated in Figure 2D-F). The acute vasodilatory effects of 17β (intact: 17.2 ± 3.7% to 50.8 ± 3.9%; abraded: 15.4 ± 2.7% to 34.4 ± 4.8%, Figure 2D) and PPT (intact: 16.7 ± 1.4% to 87.5 ± 4.0%; abraded: 24.8 ± 4.9% to 39.7 ± 8.2%, Figure 2E) were significantly blunted following endothelial abrasion. The acute vasodilatory effect of DPN did not significantly differ between groups, possibly due to the minimal responses observed to this agonist (intact: 11.4 ± 3.0% to 36.1 ± 4.6%; abraded: 16.5 ± 3.1% to 29.4 ± 5.2%, Figure 2F).

### Oestrogenic vasodilation is also impaired in non-uterine arteries from aged mice

In order to determine if the influence of ageing on oestrogenic vasodilation in uterine arteries was observable in other resistance vessels, we repeated studies on tail arteries from female mice. Figure 3 illustrates tail arterial responses to U4, ACh, 17β, PPT and DPN from 3-, 24-and 29-month-old mice. Similar to uterine arteries, both U4-induced constriction (maximum contractions for 3-month-old: 22.8 ± 1.0 kPa, 24-month-old: 18.7 ± 0.7 kPa and 29-month-old: 17.9 ± 1.0 kPa, Figure 3A) and endothelium-dependent vasodilation (maximum relaxations for 3-month-old: 57.7 ± 2.6%, 24-month-old: 43.5 ± 2.2% and 29-month-old: 40.4 ± 3.4%, Figure 3B) were attenuated in aged mice. In addition, the acute oestrogenic effects of 17β and PPT were attenuated with advanced ageing to 29 months. However, there was a trend for a decrease in the responses of tail arteries from 24-month-old mice.

**Figure 3:**
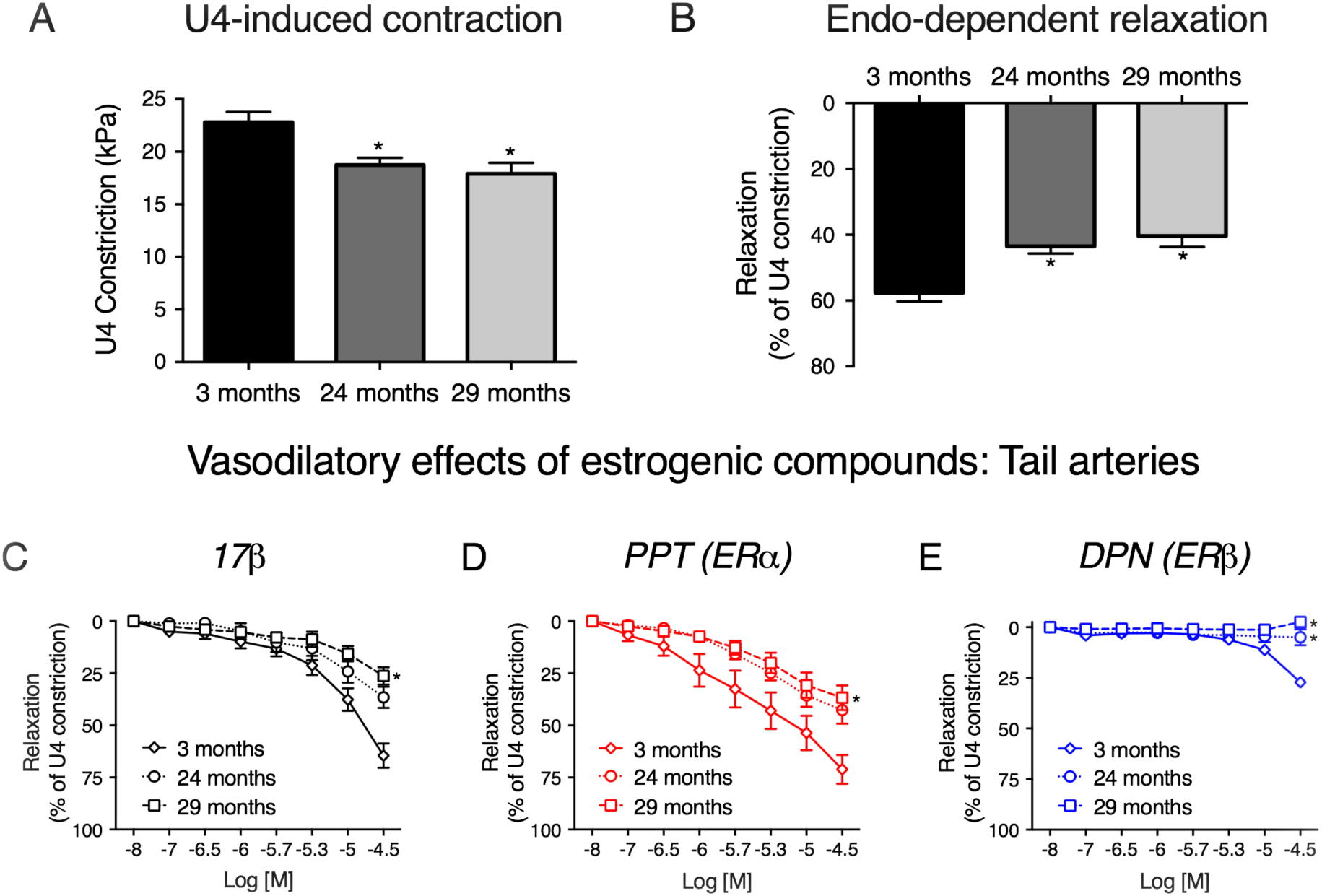
The acute responses of murine tail arteries to oestrogenic compounds are impaired by ageing. Tail arteries from 3- (n=12), 24- (n=10) and 29-month-old (n=5) mice were mounted on a wire myograph and exposed to the thromboxane agonist U4 (10^−6^M, A). Following steady-state constriction, endothelium-dependent relaxation was assessed by the addition of acetylcholine (10^−5^M, B). * P < 0.05 from young mice (unpaired *t-test*). Acute relaxations to 17β (A), PPT (B) and DPN (C) were compared in tail arteries from 3- (n=8), 24- (n=9) and 29-month-old (n=5) female mice. * P < 0.05 from young mice (repeated measures two-way ANOVA). Data are presented as mean ± SEM.

### Human arterial function and oestrogenic responsiveness is weakened in post-menopausal women

To determine whether the menopause influences vasoreactivity of resistance arteries from women, we studied the responses of uterine arteries from pre- and post-menopausal women from the ages of 32 to 75 years (Figure 4). The menopause was associated with impaired U4-induced contractility (maximum contractions for pre-menopausal: 21.7 ± 0.9 kPa and post-menopausal: 10.1 ± 0.6 kPa, Fig 4A) and endothelium-dependent relaxations (maximum relaxations for pre-menopausal: 61.5 ± 2.4% and post-menopausal: 40.7 ± 2.6%, Figure 4B) to bradykinin.

**Figure 4:**
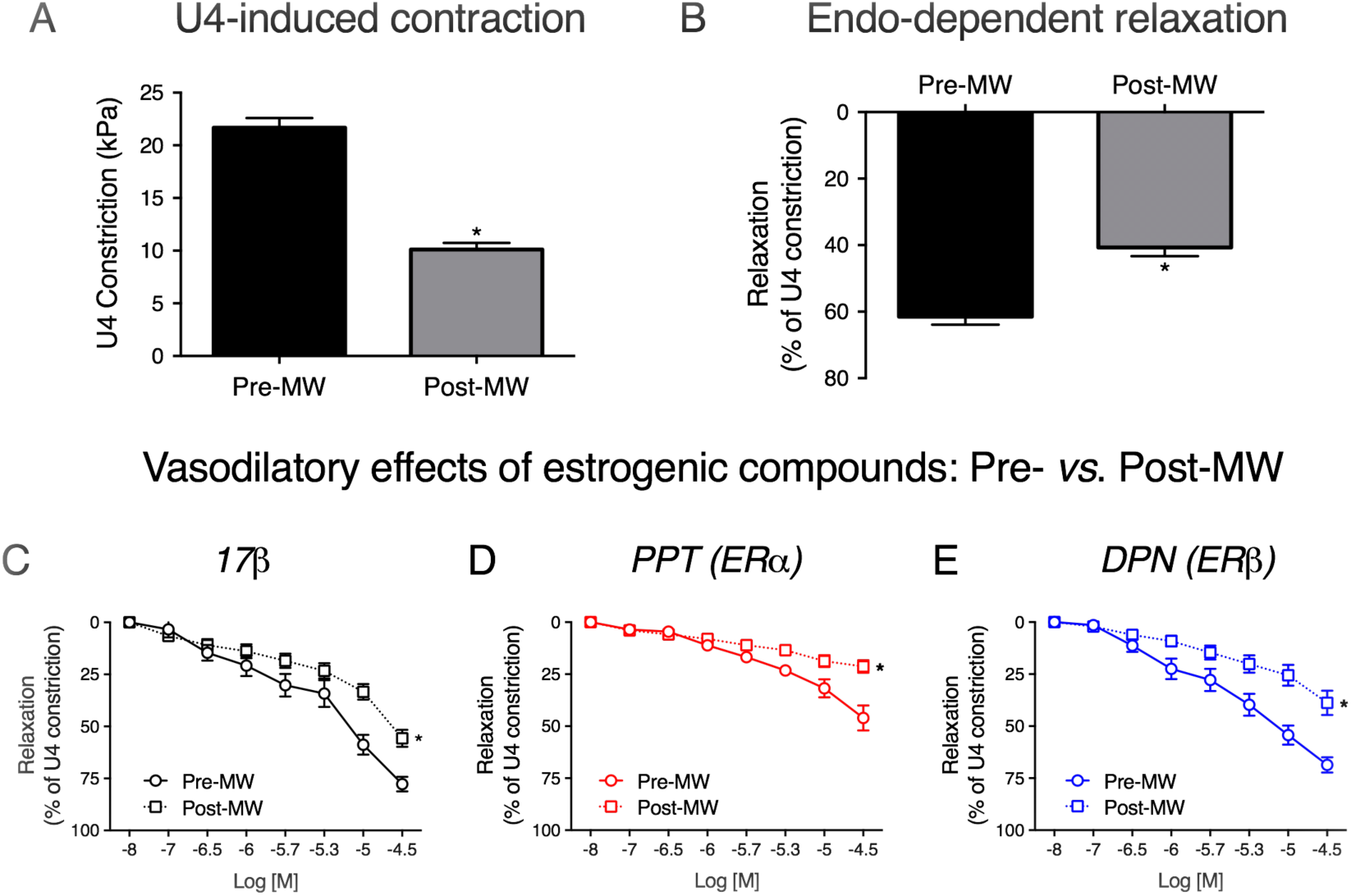
Vasoactive properties and oestrogenic responsiveness of human uterine arteries are attenuated in post-menopausal women. Uterine arteries from were mounted on a wire myograph and exposed to the thromboxane agonist U4 (10^−6^M, A). Following steady-state constriction, endothelium-dependent relaxation was assessed by the addition of bradykinin (10^−5^M, B). Contractility and endothelial-dependent relaxations were compared in arteries from pre- (n=26) and post-menopausal (n=16) women. * P < 0.05 from pre-menopausal women (unpaired *t-test*). Acute relaxations to 17β (C), PPT (D) and DPN (E) were compared in uterine arteries from pre- (n=20) and post-menopausal women (n=13). * P < 0.05 from pre-menopausal women (repeated measures two-way ANOVA). Pre-MW = pre-menopausal women. Post-MW = post-menopausal women. Data are presented as mean ± SEM.

Importantly, we found that the acute vasodilatory effects of 17β (Figure 4C), PPT (Figure 4D) and DPN (Figure 4E) were attenuated in uterine arteries from post-menopausal women.

### Human uterine arterial function is impaired with increasing age

The data presented in Figure 4 was re-plotted to examine to further examine the effect of ageing on vasoreactivity of uterine resistance arteries from women. Figure 5 demonstrates U4-induced contractions (A) and endothelial-dependent relaxations (B) in women grouped into 4 age groups (years): 30-39 (n = 7), 40-49 (n = 18), 50-59 (n = 8) and 60-75 (n = 9). Maximal U4-induced contractions were similar in women under the age of 50, but were diminished in women over 50 (maximal contractions for 30-39: 19.5 ± 1.0 kPa, 40-49: 22.6 ± 1.1 kPa, 50-59: 13.0 ± 1.0 kPa and 60-75: 8.3 ± 0.6 kPa). Interestingly, maximal endothelial-dependent relaxations were strongest in the youngest group of women and diminished in all others (maximal relaxations for 30-39: 71.8 ± 2.9%, 40-49: 57.1 ± 3.2%, 50-59: 41.4 ± 3.6% and 60-75: 41.2 ± 3.3%). Increasing age negatively influenced both smooth muscle and endothelial function (Figure 5C and D).

**Figure 5:**
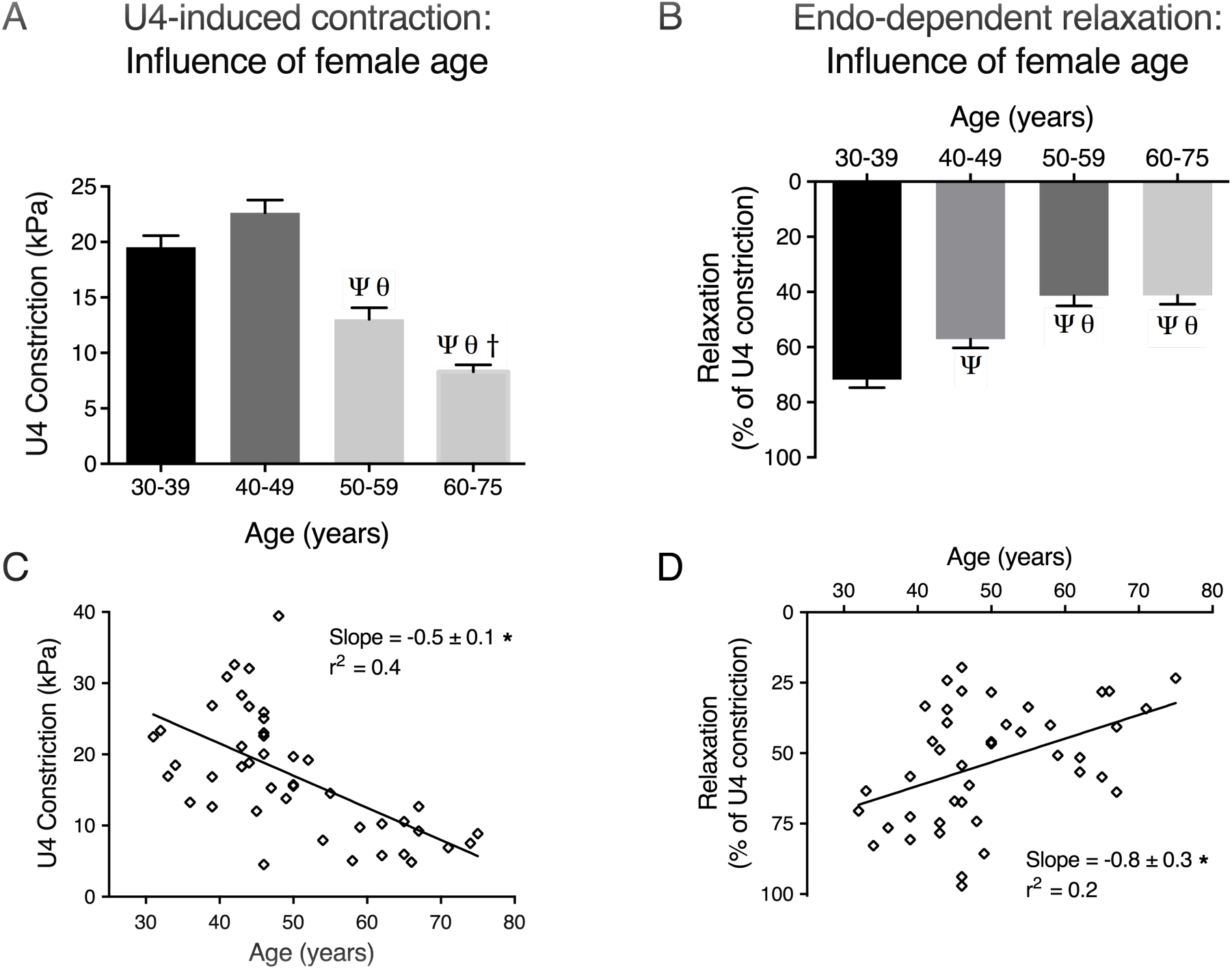
Human uterine arterial function is impaired with increasing age. Maximal U4-induced contraction (A) and endothelial-dependent relaxations (B) were compared in 4 age groups (30-39, 40-49, 50-59 and 60-75 years of age). P < 0.05 from; 30-39 (Ψ), 40-49 (θ), 50-59 (†) (ordinary one-way ANOVA). Scatter plots show (C) the mean U4-induced constriction and (D) endothelial-dependent relaxation of uterine arteries from each individual patient against plotted against age. * P < 0.05 from zero slope (*F test*). The slope and, r^2^ values are presented in each XY scatter graph.

### Oestrogenic responsiveness of human uterine arteries is impaired at older ages than changes in endothelial function

The effects of ageing on the acute responses of human uterine arteries to oestrogenic compounds were similarly assessed in female age groups (years): 30-39 (n = 5), 40-49 (n = 15), 50-59 (n = 8) and 60-75 (n = 6) (using data re-analysed from Figure 4). As shown in Figure 6A-C, the maximal relaxatory effects of 17β (maximal relaxations for 30-39: 77.1 ± 4.2%, 40-49: 78.2 ± 4.3%, 50-59: 56.8 ± 7.1% and 60-75: 58.8 ± 4.4%) and DPN (maximal relaxations for 30-39: 63.8 ± 3.8%, 40-49: 69.5 ± 5.7%, 50-59: 58.9 ± 8.5% and 60-75: 25.2 ± 4.5%) were negatively influenced by increasing age. Interestingly, whereas the vasodilatory effect of 17β was impaired in women over the age of 50, a weakening of the effects of the ER-specific agonists appeared in women over the age of 60. Similar to murine arteries, the weakened responses of human arteries to these agonists were only apparent after the occurrence of endothelial dysfunction. In addition, there was a negative trend between vasodilatory responses of human uterine arteries to oestrogenic compounds and ageing (Figure 6D&F).

**Figure 6.**
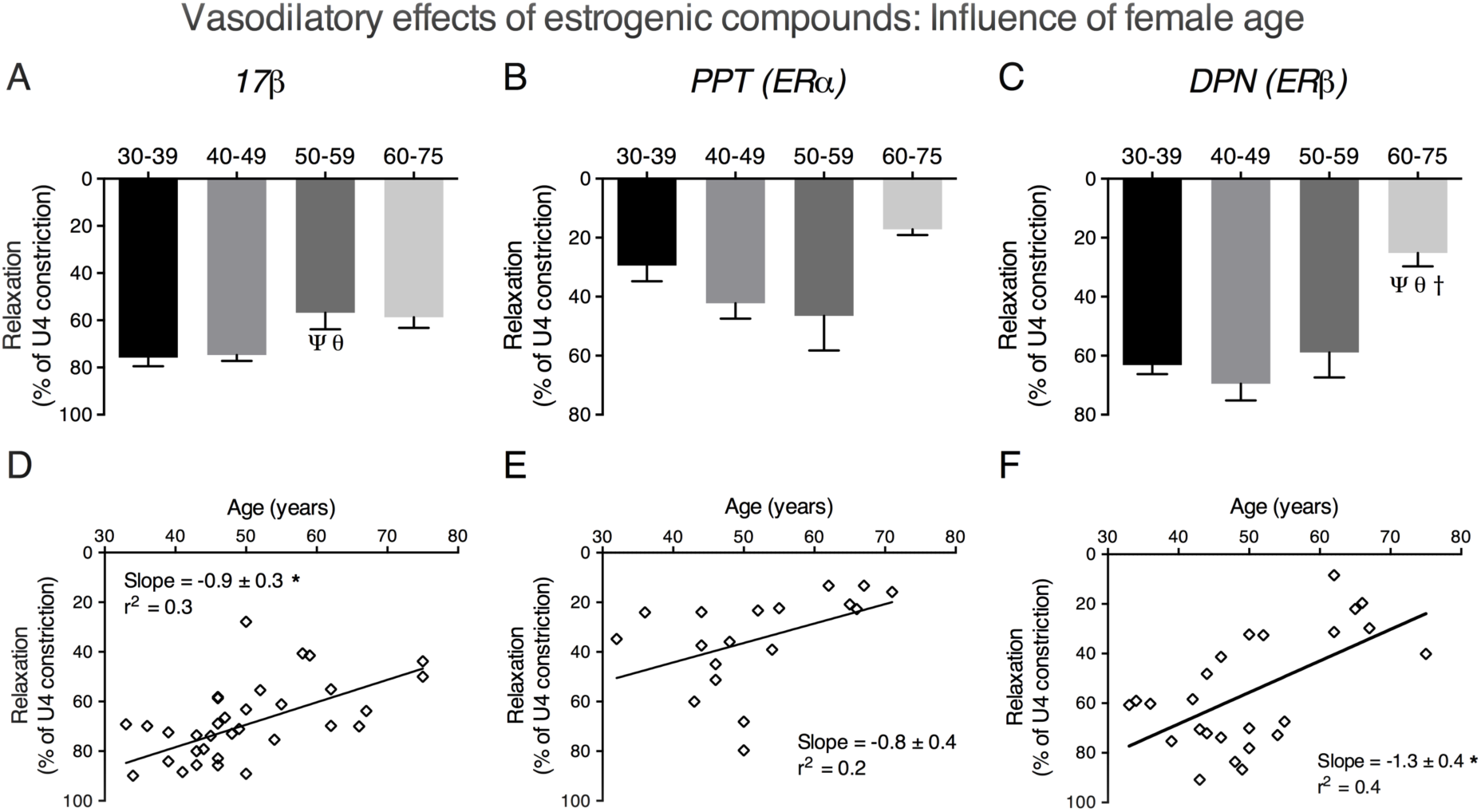
Ageing negatively regulates the acute vasodilatory effects of oestrogenic compounds in women. Maximal acute relaxations to 17β (A), PPT (B) and DPN (C) were compared in 4 age groups (30-39, 40-49, 50-59 and 60-75 years of age). P < 0.05 from; 30-39 years old (Ψ), 40-49 years old (θ), 50-59 years old (†) (ordinary one-way ANOVA). Scatter plots show the maximal relaxations from each patient plotted against increasing age. # P < 0.05 from zero slope (F test). The slope and, r^2^ values are presented at the top of each XY scatter graph.

## Discussion

The acute vasodilatory effects of oestrogen have previously been established but there is little information about how these arterial responses are influenced by the menopause and advancing age. This study investigated the influence of age and the menopause on the acute vasodilatory effects of 17β and ER-specific agonists. Our data demonstrated that both ageing and the menopause were associated with reduced vasodilatory effects of oestrogenic compounds in murine and human uterine arteries, respectively. Interestingly, however, ageing clearly altered endothelial function prior to oestrogenic responsiveness in human resistance arteries. Similarly, in mice, ageing was associated with reduced resistance arterial compliance, occurring before the altered effects of oestrogenic compounds became apparent. We therefore propose that age-related changes in arterial endothelial function and/or stiffness may be the predominating factor, rather than post-menopausal diminution of circulating oestrogen levels, which leads to impaired oestrogenic actions in arteries from post-menopausal women.

We report that the acute vasodilatory effects of oestrogenic compounds, 17β-oestradiol and ER agonists, are impaired in resistance arteries from post-menopausal women. An important consideration is what contribution ageing per se may make to these findings. Indeed, sub-group analysis of the data suggested that ageing negatively influenced the oestrogenic responsiveness of human uterine arteries. In particular, endothelial-dependent relaxations were greater in women aged 30-39 than those aged 40-49. Therefore, endothelial dysfunction of human resistance arteries occurred prior to the defect in oestrogenic responsiveness, and also the menopause. This is important when compared to the more controlled data from mice, which also demonstrated age-related impaired acute responses, in arteries from two different vascular beds, to 17β-oestradiol, PPT and DPN. Similar reductions in vascular relaxation to 17β-oestradiol have been reported for rat vessels (Lindsey et al., 2013; Wynne et al., 2004). Of note, the most pronounced effects of ageing on 17β-and PPT-induced relaxations of murine uterine arteries were observed between 24 and 30 months rather than between 3 and 24 months. This suggests that the impairment of oestrogenic responsiveness occurred after the endothelial dysfunction and altered distensibility, each evident at 24 months, were established.

Furthermore, these changes were also evident before outward hypertrophic remodeling (occurring at 29 months), suggesting changes to wall structure may be a response to worsening arterial compliance of resistance vessels.

Alterations in arterial stiffness and endothelial dysfunction are often observed in unison and the former is a strong independent indicator for subsequent cardiovascular risk (Laurent and Boutouyrie, 2015). Endothelial dysfunction with ageing has been associated with impaired nitric oxide (NO)- and EDH-mediated relaxation, each of which is a potential target of modulation by oestrogenic compounds (Wynne et al., 2004, Chan et al., 2012, Kong et al., 2015). Either way, endothelial dysfunction, or deliberate removal of endothelial-dependent relaxation, resulted in a blunting of oestrogenic responsiveness.

There is currently much debate as to the role of the menopause, and altered oestrogen bioavailability, in determining risks for cardiovascular health and attenuation in oestrogen-responsiveness of the resistance vasculature. The finding reported herein, have implications for the interpretation of clinical studies examining cardiovascular outcomes in ageing females. Certainly, oestrogen withdrawal has been reported to influence vascular gene expression, inflammation, oxidative stress, structure and function (Arenas et al., 2005; Bolego et al., 2005; Keaney et al., 1994; Kublickiene et al., 2008; Kublickiene et al., 2005; Moreau and Hildreth, 2014; Moreau et al., 2012; Pinna et al., 2008; Sudoh et al., 2001; Tarhouni et al., 2013). There is also some evidence that oestrogen therapy administered to women soon after the menopause may be of more benefit than if treatment is delayed (Giordano et al., 2015; Grodstein et al., 2006; Rossouw et al., 2007; Schierbeck et al., 2012). Recently, two clinical trials, the ‘Kronos Early Oestrogen Prevention Study’ (KEEPS) and ‘Early versus Late Intervention Trial with Oestradiol’ (ELITE) trials, have been designed to assess if the length of time since menopause alters the effectiveness of oestrogen therapy in post-menopausal women. Early data from the KEEPS trial has been disappointing, with oestrogen therapy showing no significant improvement on cardiovascular mortality, endothelial function or carotid artery compliance in young post-menopausal women (Harman, 2014; Harman et al., 2014; Kling et al., 2015). However, clinical data from the ELITE trial demonstrated that oestradiol treatment had a beneficial effect on carotid artery intima-medial thickness in recently menopausal women (< 6 years) compared with distantly menopausal women (≥ 10 years), yet secondary outcome measures of coronary atherosclerosis risk were unaffected (Hodis et al., 2016).

These outcomes necessitate a continued appraisal of the likely impact of changes in cardiovascular function related to age in women. For example, although long-term risks of adverse cardiovascular outcomes may be similar for post-menopausal women and males there is evidence suggesting key gender differences including: first cardiovascular events occurring at older ages in females and the risk of coronary artery disease less than males (Leening et al., 2014). In addition, apparent age-related gender differences in ischemic heart disease mortality in England, Wales and the United States was attributed to a deceleration of associated events in older males, rather than an acceleration in post-menopausal women (Vaidya et al., 2011). These scenarios do not fit seamlessly with a notion that the menopause per se imparts enhanced risk to female cardiovascular health.

Taking our in vitro experimental data in context with the above clinical studies, we, therefore, propose that adverse arterial ageing is a predominant factor in influencing the altered vasodilatory effectiveness of oestrogenic compounds. However, since the numbers recruited to the human study were small, we were not able to fully dissociate the effects of the menopause and ageing on the altered effects of oestrogenic compounds. Therefore, there is merit in performing a study, with larger numbers of human participants to explore further the relationship of age-related vascular dysfunction and altered oestrogenic responsiveness. This would require longer than the duration of the current study, and/or the co-operation of multiple sites. Access to other sources of resistance vasculature, such as from gluteal biopsies (Briet et al., 2013; Greenstein et al., 2010), would also extend the interpretations of findings to another human vascular bed. Such studies will be beneficial in determining the feasibility of developing new regimens of oestrogenic agents to treat vascular disease in post-menopausal women.

## Experimental procedures

### Animals

Mouse strains from the C57BL background (C57BL/Ircfa) were housed in the Comparative Biology Centre, Newcastle University, UK. Female mice from three separate age groups (3-, 24-, and 29-month-old) were utilised in the study. Mice were anaesthetised in an induction chamber using a rising concentration of 3% isoflurane/3L oxygen per min. Once anaesthesia was achieved the animal was culled by cervical dislocation, thus confirming death as required by Animals (Scientific Procedures) Act 1986. The oestrous stage of the mice was determined by vaginal smear (Evans and Long, 1922). Vaginal smears from aged mice were characterised by persistent di-oestrous as described previously (Joshi et al., 1993). Small uterine and tail arteries were obtained from mice immediately after culling and placed in ice-cold tissue collection buffer (TCB) (in mM: 154 NaCl, 5.4 KCl, 1.2 MgSO_4_, 10 MOPS, 5.5 glucose, 1.6 CaCl_2_; pH 7.4).

### Human subjects

Ethical approval was obtained from Newcastle and North Tyneside Research Ethics Committee 1 (10/H0906/71) to perform research on samples collected as part of the Uteroplacental Tissue Bank. Women who accepted to take part in the study gave written informed consent in compliance with the Helsinki Declaration. Patient characteristics are detailed in Table 2. Those who had a last menstrual period (LMP) of more than one year before the operation were deemed to be post-menopausal. Uterine arteries (identified on the myometrial surface under a dissecting stereomicroscope, at 1.5 × magnification, Leica Microsystems, UK) from pre-and post-menopausal women were obtained from lower myometrial biopsies (~1 cm^3^) taken at the time of total hysterectomy, and placed directly into ice-cold tissue TCB.

**Table 2:** Patient details. All values, apart from ethnicity and indication, are presented as mean ± SEM. Pre-MW = pre-menopausal women. Post-MW = Post-menopausal women. * P < 0.05 from pre-menopausal women (unpaired *t-test*).

| Characteristics | Pre-MW (n = 26) | Post-MW (n = 16) |
| --- | --- | --- |
| Age (years) | 42.3 ± 0.5 (33 – 50) | 61.9 ± 0.9 (50 - 75)* |
| BMI (kg/m<sup>2</sup>) | 28.3 ± 0.5 | 27.1 ± 0.4 |
| Smokers | 6 (30 %) | 2 (17 %) |
| Gravidity | 2.2 ± 0.2 | 2.5 ± 0.2 |
| Parity | 2.1 ± 0.1 | 2.0 ± 0.1 |
| Last menstrual period (years since) | - | 11.4 ± 0.8 |
| **Ethnicity:** |  |  |
| British white | 24 | 15 |
| European white |  | 1 |
| Not stated | 2 |  |
| **Indication for surgery:** |  |  |
| Menorrhagia | 13 | 1 |
| Prolapse | 3 | 12 |
| Abdominal pain | 1 |  |
| Cystocele | 2 | 1 |
| Fibroids | 3 |  |
| Prophylactic cancer reducing |  | 1 |
| Pelvic pain | 2 |  |
| Urinary symptoms | 1 |  |
| Stress incontinence | 1 |  |
| Right iliac fossa pain |  | 1 |
| **Resting diameter (µm):** | 303.2 ± 7.3 | 264.7 ± 10.4* |

### Isometric force measurements

Arteries were cut into 2-4mm segments and mounted on a small vessel wire myograph (610M; Danish Myotechnologies, Denmark) by inserting two tungsten wires (25μm for mouse arteries and 40μm for human arteries) through the lumen of the vessel. The first wire was attached to a micrometer (allowing adjustment of the lumen diameter during the normalization process) and the second wire was attached to a force transducer (which recorded changes in vessel wall tension). Vessels were normalised to a passive a passive diameter equivalent to 0.9 of L_13.3_ kPa, as described previously (Davis and Gore, 1989; Mulvany and Halpern, 1977; Mulvany and Nyborg, 1980), and equilibrated in physiological salt solution (PSS) (in mM: 127 NaCl, 4.7 KCl, 2.4 MgSO_4_, 25 NaHCO_3_, 1.18 KH_2_PO_4_, 0.07 EDTA, 1.6 CaCl_2_, 6.05 glucose bubbled with 95% air/5% CO_2_; pH 7.4) at 37°C for at least 20 minutes.

Vessels were pre-constricted with the thromboxane agonist 9,11-dideoxy-9a,11a-methanoepoxy prostaglandin F_2α_ (U4, Merck Millipore, UK) and allowed to reach a steady-state contraction. To assess endothelial function, human and mouse arteries were exposed to the endothelium-dependent vasodilators, bradykinin and acetylcholine, respectively. Arteries were then washed 3 times with PSS, pre-constricted with U4 (10^−6^M), and exposed to increasing doses (5 min duration each, of 10^−8^, 10^−7^, 10^−6.5^, 10^−6^, 10^−5.7^, 10^−5.3^, 10^−5^, 10^−4.5^M) of either 1,3,5-Estratriene-3, 17β-diol (17β-oestradiol) (Sigma-Aldrich, USA), 4,4’,4’’-(4-Propyl-[1H]-pyrazole-1,3,5-triyl)-trisphenol (PPT), 2,3-bis(4-Hydroxyphenyl)-propionitrile (DPN), (Tocris Biosciences, UK) or vehicle control (ethanol). This concentration range has been used in previous studies (Corcoran et al., 2014; Hisamoto and Bender, 2005; Montgomery et al., 2003; Scott et al., 2007; Shaw et al., 2000). In separate experiments, the endothelium of mouse uterine arteries was rendered dysfunctional by gently abrading the intimal surface with human hair. Endothelial functional integrity, determined by the relaxant effect of acetylcholine (10^−5^M) on pre-constricted arteries, was 62.4 ± 4.2% for intact and 6.4 ± 1.0% for abraded arteries indicating that abrasion had evoked endothelial vasodilatory dysfunction. Cumulative concentration response curves were then elicited with 17β, PPT, DPN and vehicle control.

Data recorded (Myodaq; Danish Myotechnologies, Denmark) as active wall tension (Δ*T* in mN/mm) was transformed to active effective pressure (Δ*T*/(diameter/2000)) denoted by kPa.

### Passive structural characteristics

Murine arteries were mounted in a pressure myography system (Living Systems Instrumentation, USA) and allowed to equilibrate for 1 hour in calcium-free PSS (PSS with no added CaCl2 plus 1mM EGTA) at 60 mmHg. Intravascular pressure was then reduced to 5 mmHg and subsequently increased to 10 mmHg, 20 mmHg and then in 20 mmHg steps to 120 mmHg. Diameters were allowed to stabilise for 5 minutes before proceeding to the next pressure step. Measurements of stable intraluminal diameter and left and right wall thickness (taken from 3 separate positions in the field of view, and averaged) were obtained at each pressure.

Measurements of intraluminal diameter and wall thickness were used to calculate the following (Izzard et al., 2006):

- Area of the lumen (μm^2^): πr^2^ (where r is the radius)
- Area of the whole artery (μm^2^): π (lumen radius + one wall thickness)^2^
- Cross-sectional area of the vessel wall (μm^2^): vessel area - lumen area
- Stress (dyne/cm_2_): Pressure (1 mmHg = 1334 dyne/cm^2^) × radius / wall thickness
- Strain (ΔD/D_0_): Change in diameter from 5 mmHg / diameter at 5 mmHg
- Elastic modulus (β): From the equation y = ae^βx^ used to fit an exponential trend-line to the stress plotted against strain.

### Statistics

All values were presented as mean ± SEM. Throughout the study, n refers to the numbers of women or mice. The passive structural characteristics of mouse arteries were compared using repeated measures two-way ANOVA (Figure 1C-G). The parameters that showed significant differences with two-way ANOVA were then analysed with a Bonferroni test for the comparison between age groups. The elastic modulus (E) was compared between murine age groups using an unpaired t-test (Figure 1H). The vasodilatory effects of increasing concentrations of oestrogenic compounds were compared using repeated measures two-way ANOVA. All maximum relaxations/contractions displayed in bar graphs were compared using an ordinary one-way ANOVA, with the exception of Figure 4, in which an unpaired t-test was used to compare arteries from pre- and post-menopausal women. For the human studies, the trend between ageing and the vasodilatory responses of oestrogenic compounds was also examined using a scatter plot. A linear regression line was plotted and significant deviation from zero slope was determined by an *F-test*. Analysis was carried out using the GraphPad Prism (6.0) software (La Jolla, CA, USA). Significance was assumed at P < 0.05.

## Acknowledgements

This research was funded by the NIHR Biomedical Research Centre (UK). We thank Julie Taggart for the processing and cataloguing of human tissue and the clinical staff and patients of Newcastle Royal Victoria Infirmary.

## Author contributions

C. J. N and M. S. performed the experiments and analysed the data. S. C. R facilitated human tissue provision. M. J. T, C. J. N and S. C. R conceived and designed the study. C. J. N and M. J. T prepared the manuscript. All authors approved the final version.

